# Single-cell analysis of clonal dynamics in direct lineage reprogramming: a combinatorial indexing method for lineage tracing

**DOI:** 10.1101/127860

**Authors:** Brent A. Biddy, Sarah E. Waye, Tao Sun, Samantha A. Morris

**Affiliations:** Department of Developmental Biology; Department of Genetics; Center of Regenerative Medicine. Washington University School of Medicine in St. Louis. 660 S. Euclid Avenue, Campus Box 8103, St. Louis, MO 63110, USA.

**Keywords:** Single-cell analysis, Direct lineage reprogramming, Clonal tracking

## Abstract

Single-cell technologies are offering unprecedented insight into complex biology, revealing the behavior of rare cell populations that are typically masked in bulk population analyses. The application of these methodologies to cell fate reprogramming holds particular promise as the manipulation of cell identity is typically inefficient, generating heterogeneous cell populations. One current limitation of single-cell approaches is that lineage relationships are lost as a result of cell processing, restricting interpretations of the data collected. Here, we present a single-cell resolution lineage-tracing approach based on the combinatorial indexing of cells, ‘CellTagging’. Application of this method, in concert with high-throughput single-cell RNA-sequencing, reveals the transcriptional dynamics of direct reprogramming from fibroblasts to induced endoderm progenitors. These analyses demonstrate that while many cells initiate reprogramming, complete silencing of fibroblast identity and transition to a progenitor-like state represents a rare event. Clonal analyses uncover a remarkable degree of heterogeneity arising from individual cells. Overall, very few cells fully reprogram to generate expanded populations with a low degree of clonal diversity. Extended culture of these engineered cells reveals an instability of the reprogrammed state and reversion to a fibroblast-like phenotype. Together, these results demonstrate the utility of our lineage-tracing approach to reveal dynamics of lineage reprogramming, and will be of broad utility in many cell biological applications.

## Introduction

Advances over the past half-century such as nuclear transfer^1^ and factor-mediated reprogramming^2^ have revealed the remarkable plasticity of cell identity. Cells reprogrammed to pluripotency can be directed to differentiate toward desired target populations by recapitulating embryonic development *in vitro,* although this approach is inefficient and produces heterogeneous populations of developmentally immature cells^3,4^. ‘Direct lineage reprogramming’ aims to directly transform cell identity between fully differentiated somatic states via the forced expression of select transcription factors (TFs). These direct strategies aim to bypass progenitor or pluripotent states^3,5^, a shortcut intended to maximize the speed and efficiency of cell fate conversion. Using this approach, fibroblasts have been reprogrammed toward many clinically valuable cell types^6–10^. It is clear, though, that many of the resulting cells do not fully recapitulate target cell identity and function^11,12^. Frequently, remnants of the starting cell fate persist^13–16^, and the cells generated appear to be developmentally immature^9,10,17^.

Unlocking the mechanisms of reprogramming will lead to improved cell fate engineering strategies. Efforts to manipulate cell identity have typically been challenging, though, due to the considerable inefficiency of current reprogramming strategies. The generation of pluripotent cells has been extensively studied and has been shown to involve two distinct stages: a long stochastic phase, and a shorter deterministic phase^18^. The stochastic nature of reprogramming initiation is likely to account for the inefficiencies observed in most cell fate conversion strategies; while many cells can initiate reprogramming, relatively few complete the process^19^. As a result, a remarkably heterogeneous population of cells emerge during reprogramming. Although studies on these populations in bulk have offered valuable mechanistic insight, it has remained a challenge to isolate rare successfully reprogrammed cells from the background noise of partially established states.

Single-cell analyses promise the resolution required to study rare events during reprogramming to pluripotency, as well as direct fate conversion between fully differentiated states^18,20,21^. The efficiency of these direct lineage reprogramming protocols typically ranges between 1-20%^2,22^. Recent technological advances to enable high-throughput single-cell RNA-sequencing (scRNA-seq), such as Drop-seq^23^ and InDrops^24^, are valuable to capture rare cells undergoing reprogramming. This is particularly advantageous for inefficient engineering protocols where as few as 1 in 100 cells represent the target species. The utility of scRNA-seq in this context has already been demonstrated: during direct reprogramming from fibroblasts to induced neurons, transition through a ‘partial’ progenitor state, that does not express classical markers was revealed^20^. Although these single-cell resolution analyses offer unprecedented insight into biological processes, there are clear limitations to the application of this technology. Collection of single-cell transcriptomes for scRNA-seq generally requires tissue disruption, resulting in the loss of spatial, temporal, and lineage relationships that are critical for thorough interpretations to be made. Several elegant computational approaches have been developed in response to these limitations; Seurat was been built, in part, to enable spatial reconstruction from scRNA-seq data^25^. In terms of temporal reordering to represent differentiation trajectories, Monocle has emerged as an extremely valuable tool, particularly for analysis of *in vitro* cell reprogramming and differentiation^21,26^.

Presently, many aspects of cellular behavior observed and assessed via these single-cell analytical approaches are inferred, particularly with respect to the order of events during reprogramming. This could be especially problematic in protocols involving the generation of many potential branch points. Thus, preserving information on lineage relationships between cells is critical in order to precisely map the trajectories leading to successful reprogramming outcomes. Here, we present a methodology, ‘CellTagging’, to enable combinatorial indexing of cells and readout of lineage information at the transcript level, concomitant with single-cell transcriptome analysis. This is a simple lentiviral-based tool that can be easily applied to the study of many cellular reprogramming or differentiation strategies. Here, we apply this method to the high-throughput single-cell analysis of fibroblast to induced endoderm progenitor (iEP) conversion, a valuable self-renewing engineered cell type that has the potential to functionally engraft liver^10^ and intestine^13^. iEP generation represents a prototypical direct lineage reprogramming methodology that reflects conversion via a progenitor-like state^11,13,27^, an important emerging theme in direct reprogramming^12,20^. Via this approach we have begun to map the transcriptional landscape of this reprogramming strategy, revealing potential roles for Igf2 and Notch signaling in the process. We find that while reprogramming is initiated in many cells, few reach a fully-reprogrammed state. On the clonal level there is a tremendous amount of heterogeneity, and successful reprogramming is not accounted for by distinct ‘elite’ cells that are predisposed to successfully reprogram. We also find that the emerging population of iEPs is generated from a smaller founder population, of which a dominant clone rapidly expands, presenting implications for the properties of the engineered cells. Finally, far from representing a homogenous collection of cells following this clonal expansion period, we present evidence that the population is in dynamic flux between several distinct transcription states.

## Results

### Single-cell analysis of direct lineage reprogramming from fibroblasts to induced endoderm progenitors

We previously developed a computational platform, ‘CellNet’, to evaluate cell identity via gene regulatory network reconstruction^14^. Via this approach, we found that many cell engineering protocols produced partially converted and developmentally immature cells^13,14^. We applied CellNet analysis to a direct lineage reprogramming protocol that originally aimed to convert fibroblasts directly to hepatocytes via forced expression of the endoderm transcription factors, Foxa1/2/3, and Hnf4a^10^ (**Fig. 1A**). Analysis of the resultant cells revealed a failure to silence the fibroblast expression program, weak establishment of hepatic identity, and the unexpected emergence of intestinal identity^13^. Ultimately, these cells were able to functionally engraft a mouse model of induced colitis, fully differentiating to mature intestine^13^. Given that these cells have also been shown to possess hepatic potential^10^, and self-renew *in vitro*^10,13^, we re-designated these cells as ‘induced endoderm progenitors’ (iEPs)^13^.

**Figure 1.**
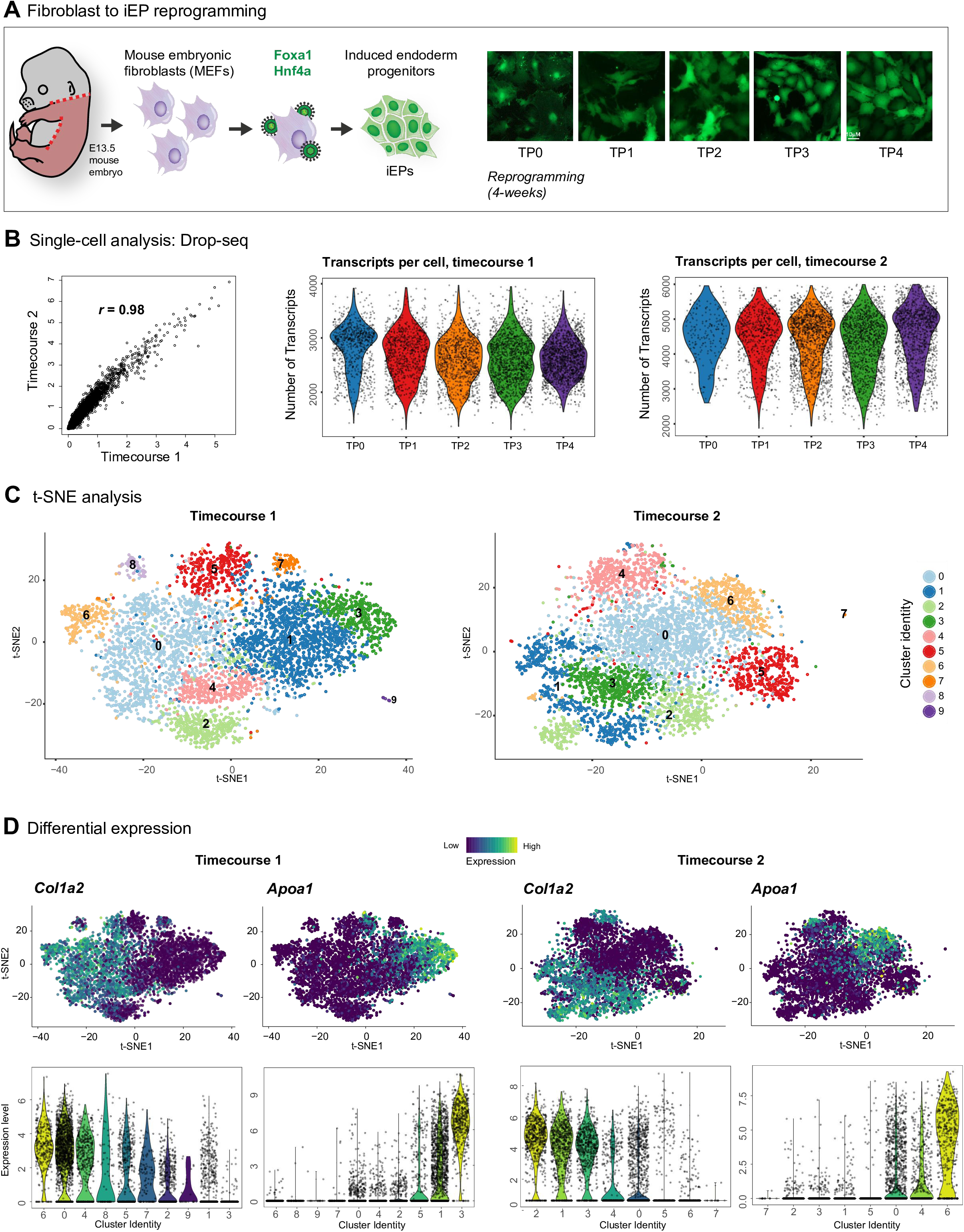
Single-cell analysis of fibroblast to iEP direct reprogramming. **(A)** Experimental overview: E13.5 mouse embryonic fibroblasts were directly reprogrammed to iEPs via retroviral delivery of Foxa1 and Hnf4a, as previously reported^10,13^. At weekly intervals throughout four-weeks of reprogramming, cells were collected for Drop-seq analysis. Non-reprogrammed fibroblasts were also harvested. **(B)** Single-cell analysis metrics. Left-panel: Scatter plot of average gene expression values (across cells) between both experimental timecourses. Correlation between these two experimental replicates is measured using the Pearson correlation coefficient. Right panels: Violin plots of UMIs (transcripts) per cell for both timecourse replicates. **(C)** Visualization of single-cell clusters using 2D t-SNE (Seurat). 10 and 8 transcriptionally distinct clusters of cells were detected in biological replicate timecourses 1 and 2, respectively. **(D)** Visualization of key changes in gene expression. Expression levels of the fibroblast marker, *Col1a2*, and the iEP marker, *Apoa1*, were projected onto t-SNE plots (upper panels) to broadly classify distinct clusters of cells. *Col1a2* is downregulated in reprogrammed cells (positioned to the right) which gradually upregulate *Apoa1* expression. Expression changes are also shown on violin plots (lower panels).

In this study, we directly reprogrammed mouse embryonic fibroblasts to iEPs via expression of Foxa1 and Hnf4a, over two independent timecourses. Two weeks after initiation of reprogramming, iEPs emerge as phenotypically distinct colonies of small cells (**Fig. 1A**). Every week, over a four week period, we harvested cells for high-throughput scRNA-seq via Drop-seq^28^. Drop-seq is a cost-effective scRNA-seq platform, employing microparticle beads coated with poly-thymidine-tagged oligonucleotides. Each oligonucleotide possesses a 12-base pair (bp) cell barcode, shared across all sequences on the same bead, and an 8-bp unique molecular identifier (UMI) that labels each captured mRNA molecule. Using a microfluidic device, beads in lysis buffer are co-encapsulated with cells. Upon cell lysis, polyadenylated mRNA transcripts are hybridized to the poly-thymidine oligonucleotides. This labels all mRNA molecules in a droplet with a cellular barcode and an individual UMI for downstream computational analysis. Following recovery, cDNA amplification generates a library of 3’ transcript ends tagged with a barcode denoting cell-of-origin. We performed a species-mixing experiment to confirm single-cell resolution capture in our hands (**Fig. S1A**).

At each stage of reprogramming, including the original fibroblast population, we harvested 150,000 cells for Drop-seq, aiming to capture 5000 single-cell transcriptomes. Following library preparation and sequencing, according to^28^, processed reads were mapped to the mm10 mouse genome assembly using STAR and digital gene expression matrices were generated using the Drop-seq tools pipeline (http://mccarrolllab.com/dropseq/). Cells expressing 200 or more genes were selected for inclusion in the matrices. Over both timecourse experiments, this resulted in a total of 18,737 cells with mean counts of 1,517 genes and 4,811 UMIs/transcripts per cell. Averaged expression levels of genes were highly correlated between the two biological replicates (**Fig. 1B**). We subsequently filtered these matrices to include only those cells with a UMI count of at least 1000, discarding cells in which the proportion of the UMI count attributable to mitochondrial genes was less than or equal to 10%. This yielded a total of 11,346 cells with mean counts of 1,810 genes and 6,163 UMIs/transcripts per cell for downstream analysis (**Fig. 1B**).

In order to broadly assess this direct reprogramming protocol, we used a combination of the R packages Scater^29^ and Seurat^25^ to calculate cell cycle phase, normalize gene expression, and visualize clusters of transcriptionally similar cells (**Fig.S1B**). Unsupervised identification of highly variable gene expression was employed for dimension reduction via principal component analysis (PCA), where the resulting PCs were used as input to cluster the cells, using a graph-based approach. t-Distributed Stochastic Neighbor Embedding (t-SNE)^25^ was used to separate individual cells in multidimensional space and visualize them onto a 2D-map. t-SNE resolves the reprogramming process into 8-10 clusters of transcriptionally distinct cells (**Fig. 1C**). By projecting expression of the fibroblast-related gene, *Col1a2*, onto these plots, we were able to locate unconverted cells. Likewise, visualization of a gene coupled with iEP emergence, *Apoa1*, assisted in the identification of clusters harboring reprogramming cells (**Fig. 1D**). Surprisingly, considering the low efficiency of reprogramming, a high proportion of cells (∼30%) express *Apoa1*. These transcriptional changes were consistent across the two biological replicates.

### Benchmarking single-cell analyses to reveal transcriptional changes over the course of reprogramming

Single-cell analyses of biological phenomena that involve gradual transcriptional changes over time are complicated by the lack of means to precisely identify cell types and transitions. In addition, we face challenges to identify the transcriptional changes that occur due to reprogramming rather than the aging of cells in culture. Furthermore, capture of cells at different experimental intervals can introduce technical variation, potentially masking true biological variation. In an effort to overcome these limitations, we developed a multiplexing approach, spiking-in a defined population of cells to act as an internal control in each experiment. In this instance, we employed unconverted fibroblasts cultured in parallel to reprogramming cells, thus acting as an age-matched ‘benchmark’. To label these cells with an index to permit their subsequent identification, we transduce benchmark fibroblasts with lentivirus carrying GFP and a SV40 polyadenylation signal sequence. Contained within the GFP UTR is a defined 8nt index sequence. This design results in the generation of abundant, indexed and polyadenylated transcripts that are captured as part of the standard Drop-seq library preparation (**Fig.2A**). In order to maximize detection of these index sequences, we have developed a PCR method to amplify the GFP UTR harboring the index. We spike this amplicon into the cDNA library. Following sequencing, a large number of GFP-index reads are detected, comparable to the most abundant cellular transcripts (**Fig.S2A**).

**Figure 2.**
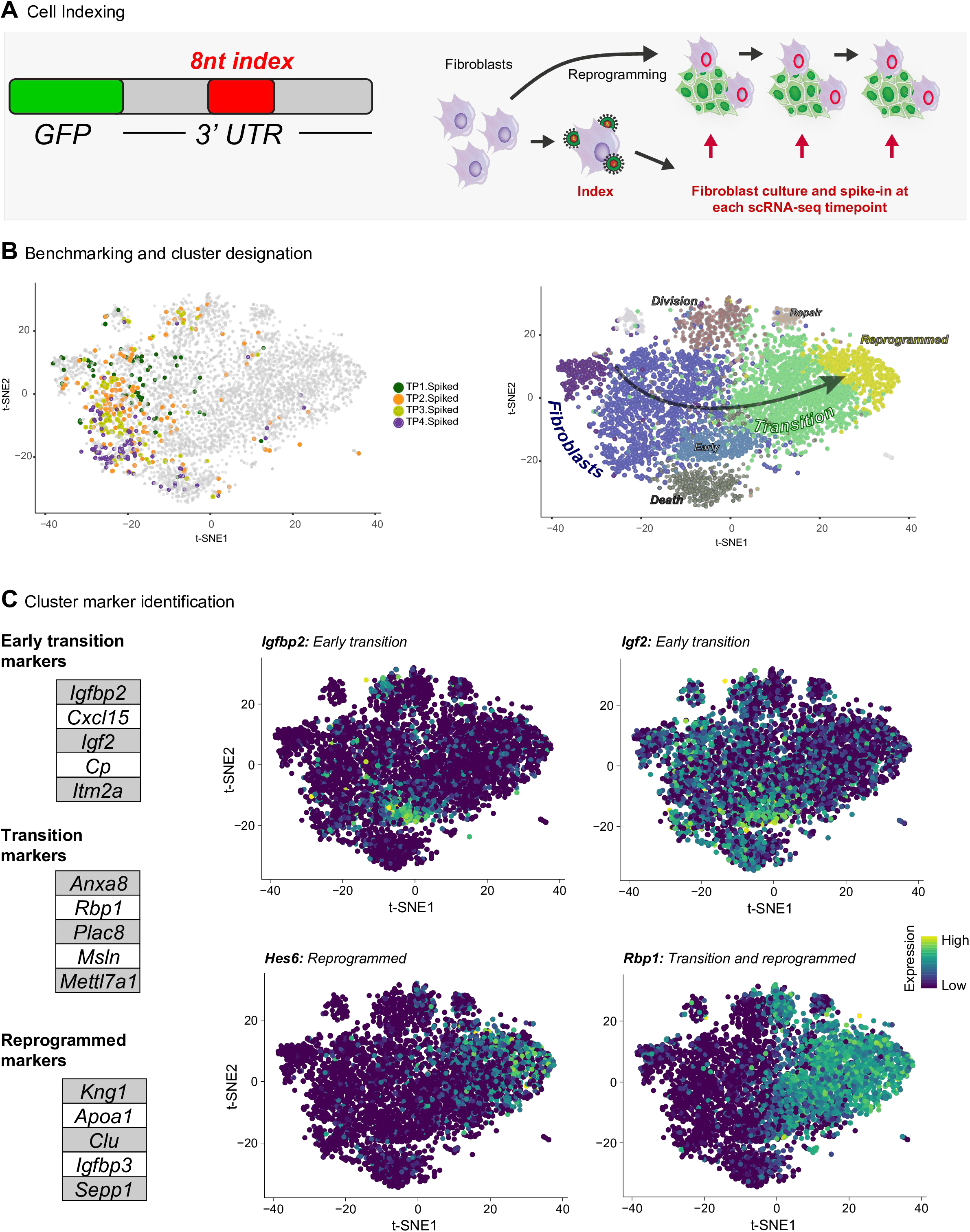
Benchmarking single-cell analyses to reveal reprogramming-related transcriptional changes. **(A)** Cell Indexing strategy: benchmark, age-matched fibroblasts were labeled with a specific 8nt index sequence embedded in the 3’UTR of GFP, delivered via a lentiviral construct. Following labeling, benchmark fibroblasts were cultured in parallel to reprogramming cells and spiked into each cell population harvested for Drop-seq analysis. These cells were spiked-in at a proportion of 10-20% of the overall population and act as an internal control for each reprogramming timepoint. **(B)** Left panel: Identification and visualization of benchmark fibroblasts. The location of these control fibroblasts is marked within 2D on the t-SNE plot for timecourse 1, where they generally cluster together. Right panel: designation of cluster identity. Each cluster on the t-SNE plot was assigned a phase of reprogramming or behavior, based on a combination of differential gene expression and gene ontology analysis. The population comprises of 7 defined clusters: 2 fibroblast clusters, early-transition, transition, reprogrammed, death, cycling, and repair. **(D)** Identification and visualization of gene expression defining specific clusters. Left tables: gene expression significantly enriched in early transition, transition, and reprogrammed stages. Top right panels: visualization of the early transition markers, *Igfbp2* and *Igf2*, using t-SNE. Bottom panels: visualization of *Hes6*, marking the reprogrammed cluster, and *Rbp1*, marking both the transition and reprogrammed clusters. All analysis here is shown for timecourse 1. Analyses for timecourse2 can be found in the supplementary material.

We implemented this approach by spiking benchmark fibroblasts into the reprogramming population at each experimental timepoint. A demultiplexing step was added to the sequencing analysis in order to identify the benchmark fibroblasts, representing 10-20% of the total population. The locations of these cells were then mapped onto t-SNE plots (**Fig.2B;S2B**). The majority of benchmark fibroblasts are located within two distinct clusters, enabling designation of these sub-populations as unconverted fibroblasts (**Fig.2B: Right panel**). Immediately adjacent to the largest of these clusters, lies a smaller group of cells, devoid of benchmarking fibroblasts, suggesting that cells within this cluster are in the first phases of reprogramming. Thus, we designated this group an ‘early-transition’ cluster (**Fig.2B**). Differential gene expression analysis reveals markers defining this transcriptional state, including upregulation of Insulin-Like Growth Factor 2 (*Igf2*) and Insulin-Like Growth Factor Binding Protein 2 (*Igfbp2*) expression (**Fig.2C**). Some of the first transcriptional changes are seen within this transition cluster, although fibroblast-associated gene silencing is not yet observed at this stage (**Fig.1D**). See **Table 1** for significant differential gene expression defining each cluster.

**Table 1.**
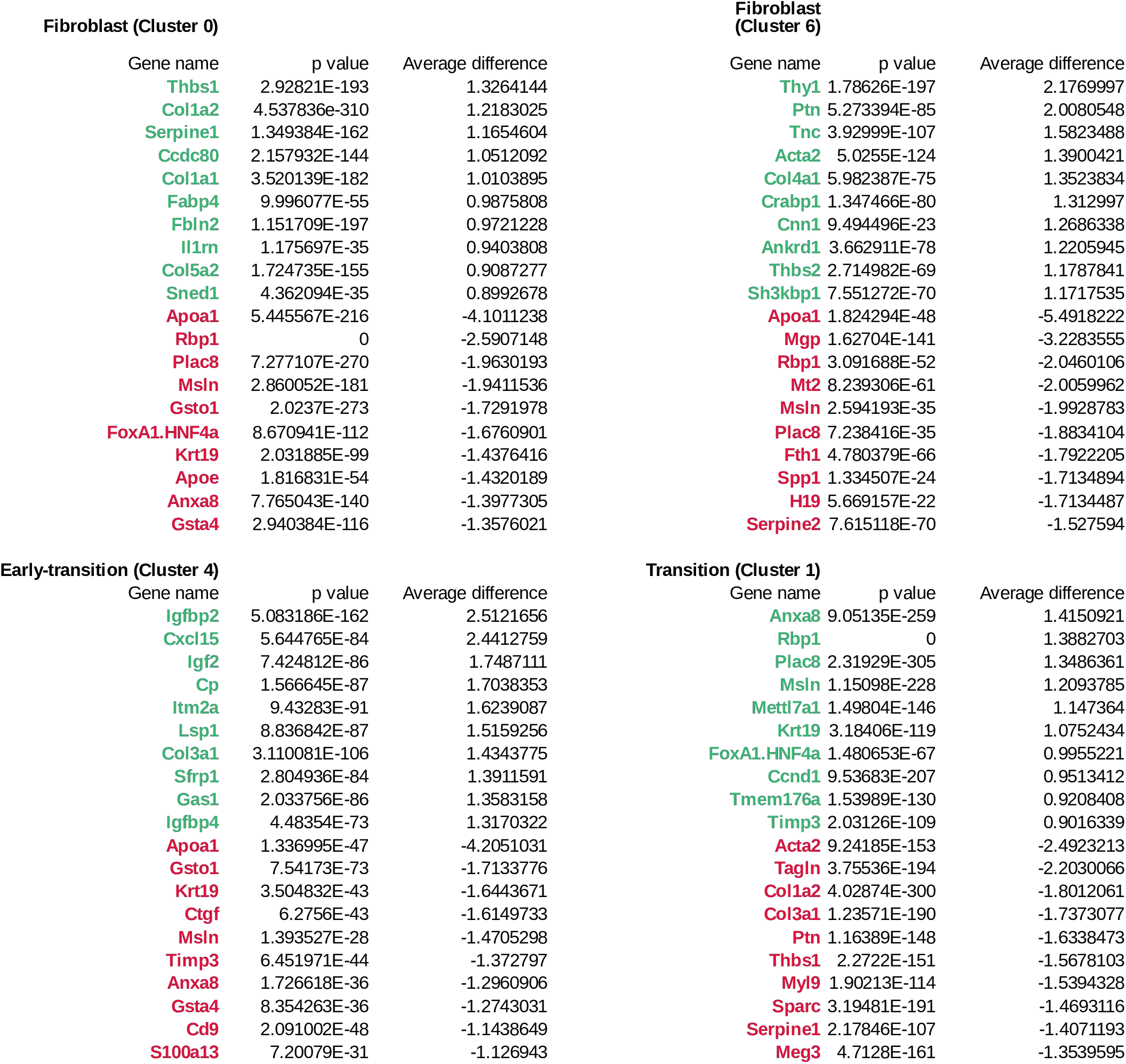
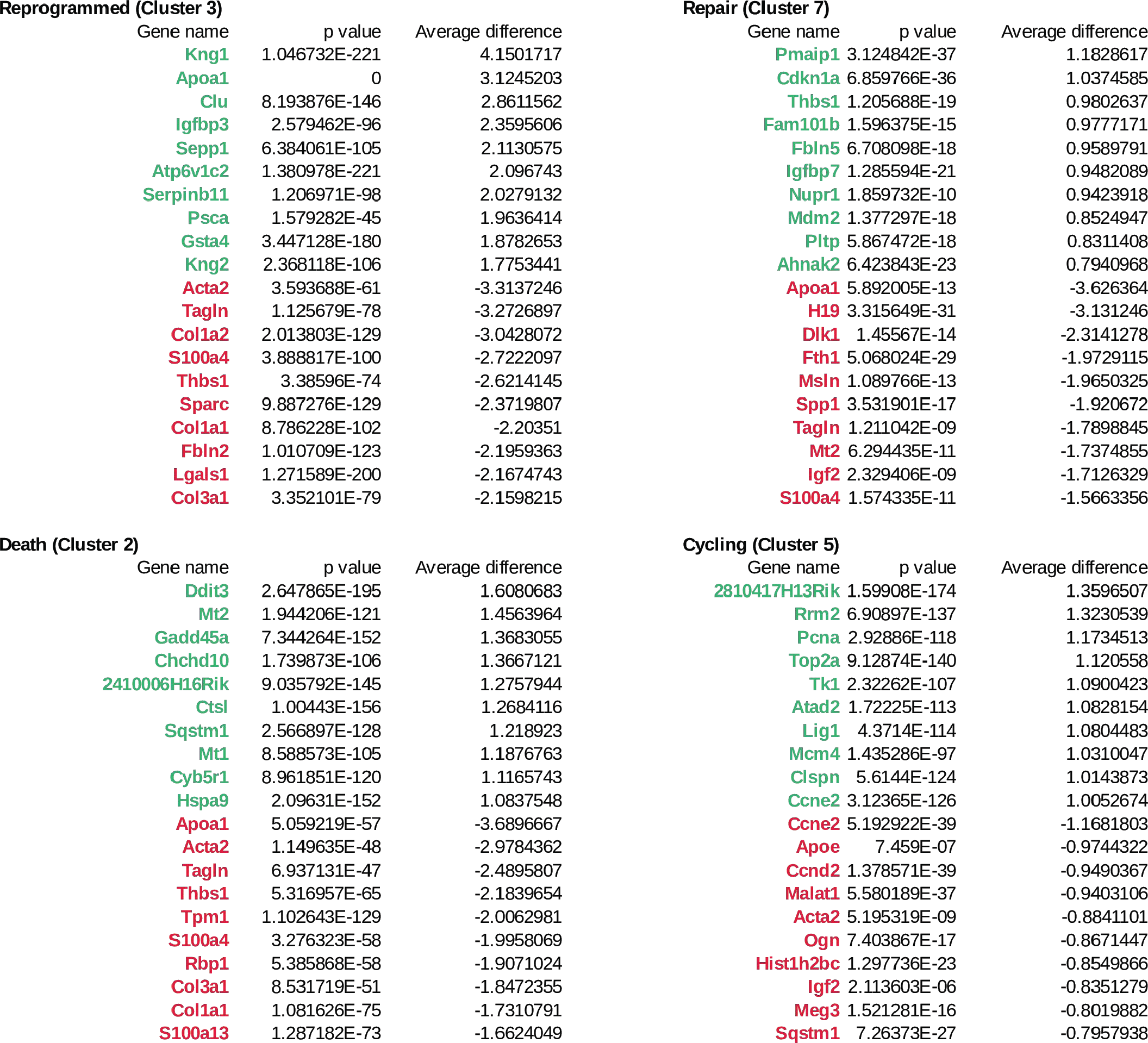
Identification of marker genes for each cluster identified for timecourse 1. Differential gene expression analysis identifies signatures of cell clusters in timecourse 1. The top 5 enriched genes marking each cluster are shown in green. The top 5 depleted genes for each cluster are shown in red.

Silencing of fibroblast gene expression is first observed in a large cluster of cells (**Fig1C**, cluster 1: timecourse 1, cluster 0: timecourse 2), accompanied by the initiation of genes expressed in iEP populations^10,13^. We named this group the ‘transition’ cluster (**Fig.2B;C**). Notably, Retinol Binding Protein 1 (*Rbp1*) marks cells within this transition state. Rbp1, plays a role in the transport of vitamin A, a molecule that has recently been shown to play a role in the erasure of epigenetic memory during reprogramming to pluripotency^30^. This primary group of clusters concludes with putative ‘reprogrammed’ cells, expressing the highest levels of iEP-associated genes. These cells also express high levels of *Hes6*, a TF component of the Notch signaling pathway^31^ (**Fig.2C;S2C**). In this reprogrammed cluster, Albumin (*Alb*) expressing-cells are rare (data not shown), supporting earlier reports of weak hepatic identity in these cells^13^. Other smaller clusters are defined as dying, cycling, and cells undergoing DNA repair, based on GO annotations (**Fig.2B**).

Altogether, these analyses help position single-cells within discrete phases of the reprogramming process. A high proportion (∼30%) of cells appear to initiate reprogramming, although previous colony formation assays estimate that only 1-2% of cells become fully reprogrammed to iEPs^10,13^. This discrepancy indicates that many cells are partially reprogrammed, a state that has previously been described during reprogramming to pluripotency^2,18^. These partially reprogrammed cells may be lost via apoptosis before they reach a fully reprogrammed state. This notion is supported by previous studies demonstrating cell death as a factor that reduces the efficiency of reprogramming to pluripotency^32^. To explore these possibilities, we require a more accurate method to differentiate between fully-and partially-reprogrammed cells within this larger population of single cells.

### Scoring cell identity based on bulk expression distinguishes between partially-and fully-reprogrammed cells

From the above differential expression analyses, we were unable to identify gene expression that exclusively marks the putative reprogrammed cell cluster. To resolve this, we adapted an approach that was previously used to score cell identity during direct reprogramming from fibroblasts to induced neurons^20^. In this method, quadratic programming is employed to deconstruct each single-cell transcriptome and represent its identity as a fraction of the starting and target cell types. We have adapted this approach, using the KeyGenes^33^ platform to curate a list of genes defining fibroblast and iEP cell identity, based on the original microarray profiling from Sekiya and Suzuki^10^. **Fig.3A** shows fibroblast and iEP fractional identities for all cells in timecourse 1, ordered according to loss of fibroblast identity, with concomitant establishment of iEP identity. The majority of cells retain considerable fibroblast identity, defined as a fractional fibroblast score of >0.8. Around 2% of cells are classified as fully reprogrammed, receiving a fractional iEP score of >0.8, whereas 10% of cells are within a transition between these two states. This is also reflected in the second timecourse (**Fig.S3A**). This rarity of fully reprogrammed cells is in agreement with earlier efforts to measure efficiency of this reprogramming protocol via colony formation assays^10,13^. To validate the utility of this approach, we used the Monocle2 package^26^ to reconstruct direct reprogramming in pseudotime. Monocle uses dimension reduction to represent each single-cell in 2D space and effectively ‘joins-the-dots’ to construct a differentiation trajectory, a ‘pseudotemporal’ ordering of cells based on the gradual re-wiring of their transcriptomes. Projection of fractional iEP identity onto this trajectory shows accordance between these two computational approaches to assess the reprogramming process (**Fig.3A;S3A, right panels**). Moreover, visualization of fractional fibroblast and iEP scores onto t-SNE plots indeed confirms that the fully reprogrammed cells are harbored into our predicted reprogrammed cell cluster, with diminished fractional fibroblast identity scores appearing in what we had defined as the transition cluster (**Fig.3B;S3B**). We used this approach to more accurately detect iEP emergence, and found that fully-reprogrammed cells emerge by 2-weeks post-initiation of reprogramming. In the first timecourse, the number of iEPs peak at week 3, and week four in the second timecourse (**Fig.3C;S3C**). Using *Apoa1* expression as a marker of reprogramming initiation, we overlayed these expression values onto iEP fractional identity scores (**Fig.3D**). This revealed that many cells (∼30%) appear to initiate reprogramming, whereas projection of fibroblast-specific *Col1a2* expression indicates that fibroblast gene expression fails to be silenced in many of these cells (**Fig.S3D**). We next wanted to explore these distinct reprogramming trajectories, to examine the origins of reprogrammed cells, and the eventual fate of partially reprogrammed cells. This is challenging given the asynchronicity of reprogramming, leading to a great deal of heterogeneity at each experimental timepoint. Thus, we aimed to reduce the complexity of these analyses by tracking individual clones of cells undergoing reprogramming.

**Figure 3.**
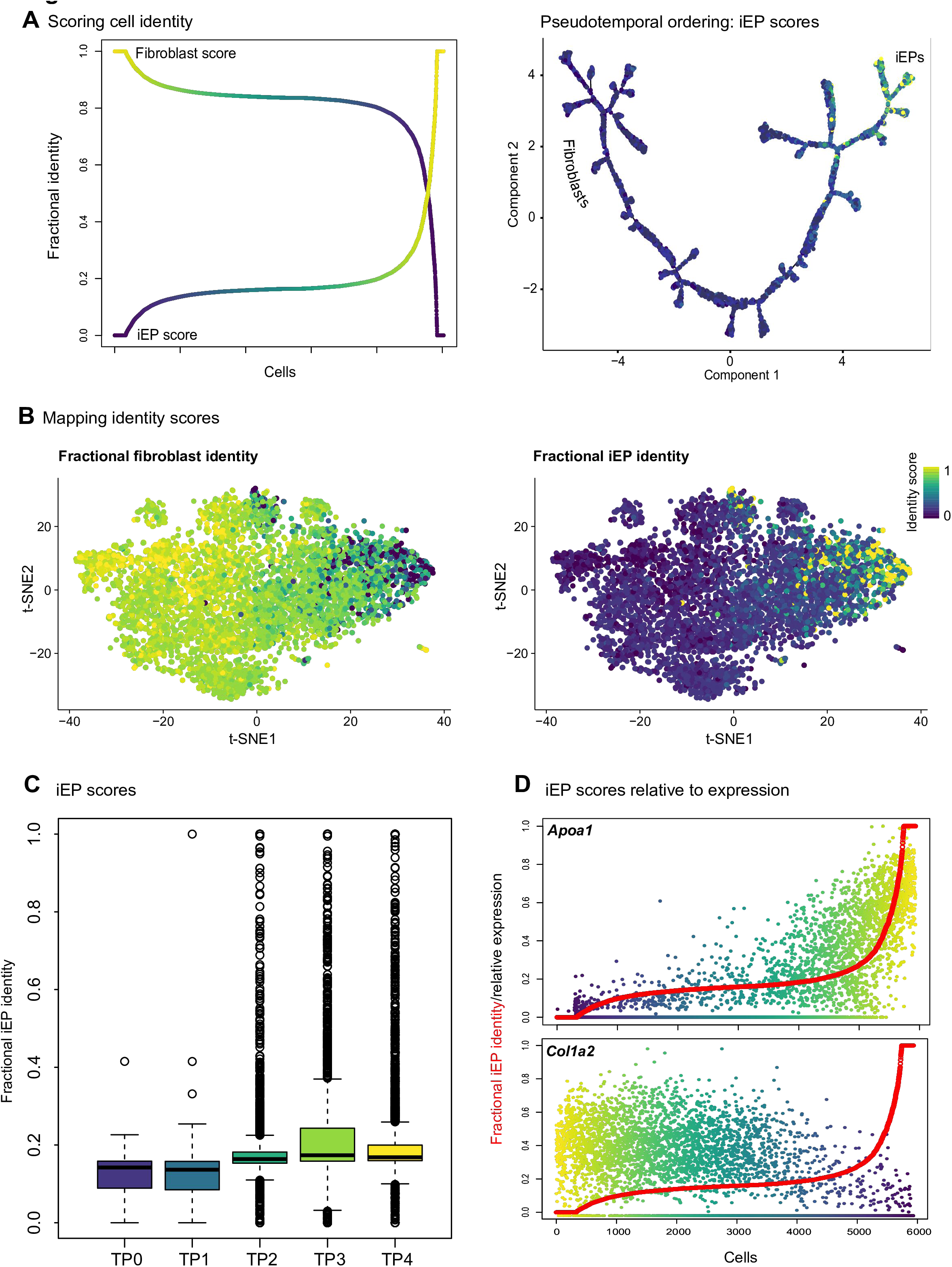
Scoring cell identity using quadratic programming. **(A)** Left panel: For all timecourse 1 cells, similarity to the bulk transcriptome for fibroblasts and iEPs was calculated using quadratic programming and plotted as fractional identities. Cells were ordered according to increase in iEP identity. Right panel: fractional iEP identity scores projected onto Monocle reconstruction of the reprogramming in timecourse 1. The end of reprogramming, as defined by Monocle, coincides with high iEP scores. **(B)** Projection of fibroblast (left panel) and iEP (right panel) fractional identity scores onto t-SNE visualizations of cell clusters. Emergence of iEPs overlaps with transition and reprogramming clusters in this first timecourse. **(C)** Boxplot of iEP fractional identity scores, grouped by timepoint. **(D)** Plot showing relative expression of Apoa1 (top panel) and Col1a2 (lower panel) with fractional iEP scores overlayed (red). Cells are according to increase in iEP identity.

### CellTagging: a combinatorial indexing method to track clonal dynamics of direct reprogramming

In current single-cell transcriptome analyses, information on lineage relationships between cells is lost. This knowledge is essential for tracing the fate of partially-and fully-reprogrammed cells. Several elegant lineage tracing solutions such as the Sleeping Beauty transposase^34^ and a CRISPR/Cas9 self-evolving barcode^35^ exist but are not presently compatible with scRNA-seq. We aimed to simultaneously read-out the single-cell transcriptome along with lineage information revealing the reprogramming history for each cell. To this end, we have developed a lineage-tracing approach to reveal clonal ancestry in parallel with cell identity. Briefly, each individual cell in the original fibroblast population is labeled with a unique combination of genetic DNA indexes, ‘CellTags’, introduced via lentiviral transduction (**Fig.4A**). We based this methodology on our approach to index benchmark fibroblasts, modifying the approach to engineer random 8nt sequences into the 3’UTR in place of the defined index sequence. We generated a complex library where each lentivirus could carry one of 64,000 unique CellTags. We transduced fibroblasts with this library at a multiplicity of infection ∼10, resulting in each fibroblast being labeled with a unique combination of ∼10 CellTags (**Fig.S4A**). Reprogramming to iEPs was initiated 24hr post-CellTagging. We again employed our PCR amplification method to maximize sequencing coverage of CellTags, ensuring reliable detection in parallel with scRNA-seq readout (**Fig.S2A**).

**Figure 4.**
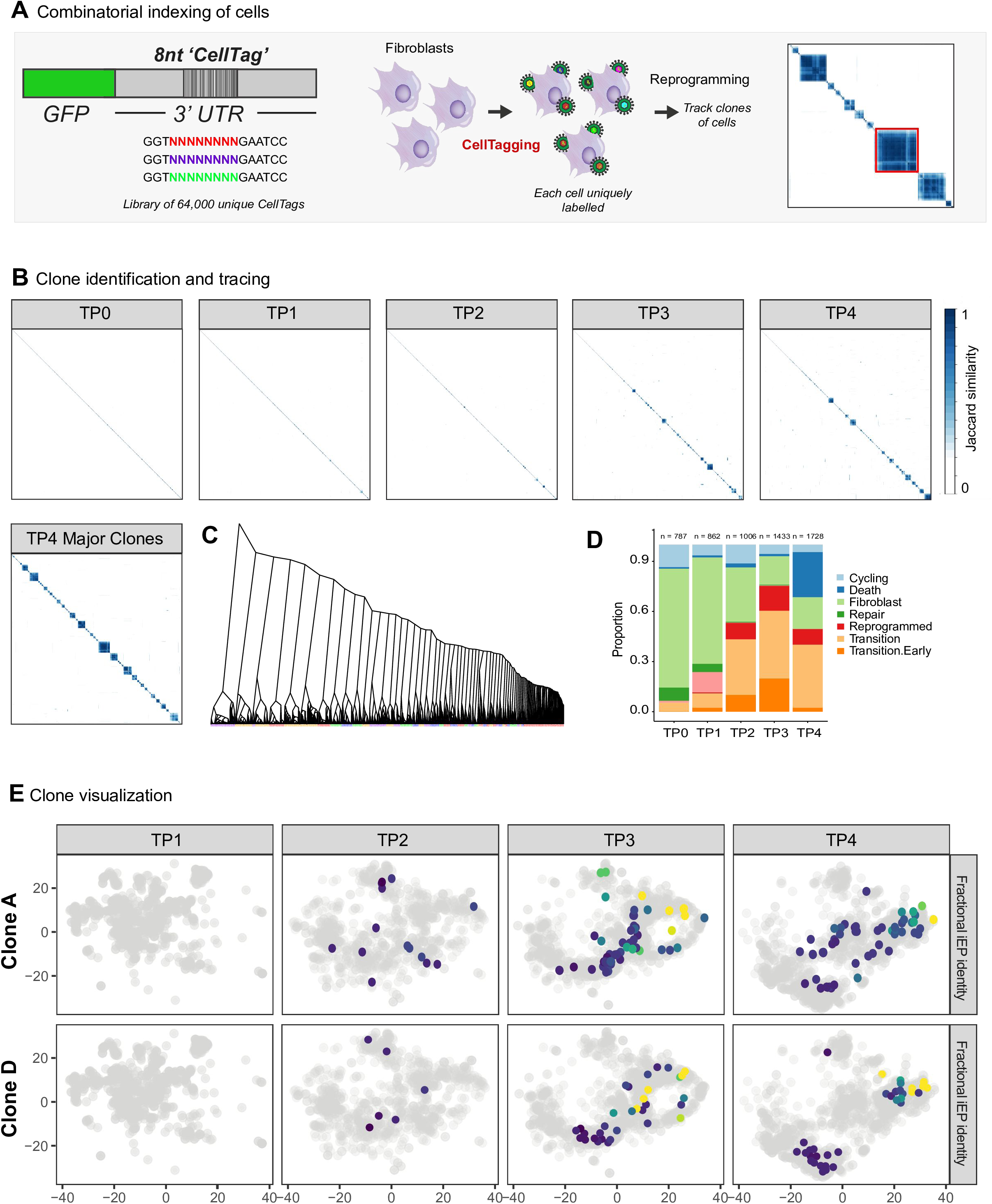
CellTagging: a combinatorial labeling strategy to track clonal relationships between cells. **(A)** The CellTagging methodology. We adapted our cell indexing and benchmarking approach by replacing the defined 8nt index in the 3’UTR of GFP with an 8nt random sequence (CellTag). This resulted in a complex library of up to 64,000 unique CellTags. Lentivirus was prepared with this library and used to transduce fibroblasts at a multiplicity of infection ∼10, resulting in the labeling of each fibroblast with a unique combination of CellTags. Fibroblasts were then cultured as benchmark cells or reprogrammed to iEPs. At each experimental timepoint, CellTag data was collected in concert with each single-cell transcriptome and unique CellTag signatures were extracted for each cell. Weighted Jaccard analysis was used to calculate the similarity of CellTag signatures between cells and is presented as correlation plots. Hierarchical clustering is used to group clones of cells (based on CellTag signature overlap) together on these plots. Right panel: the red outline demonstrates the identification of a clone of cells derived from the same initial CellTagged fibroblast. **(B)** Clone identification and tracking over multiple timepoints. CellTag signatures were extracted for all cells in timecourse 1 and are presented as correlation plots. Upper panels: Over the course of reprogramming, distinct clones emerge. Lower panel: correlation plot of major clones identified in timepoint 4. **(C)** Dendrogram showing relationships between cells. This analysis was used to identify distinct clones of cells, some of which could be tracked over multiple timepoints. **(D)** Cell identity/behavior designation for all cells over the course of reprogramming in timecourse 1. This is used as a point of reference for individual clone behavior. **(E)** Behavior of two representative clones. Cells belonging to clone A are detected from timepoints 2 to 4 and are detected in all stages of the reprogramming process, demonstrating a high degree of heterogeneity. Cells of clone D are also detected from timepoints 2 to 4. Many cells of this clone are found in the transition cluster at timepoint 3. At timepoint 4, this has resolved into two distinct outcomes: successful reprogramming, or death.

Clonally-related cells were identified via overlap of their combinatorial CellTag signatures. To recover these sequences, reads were extracted using the CellTag “motif” (GGTNNNNNNNNGAATTC). These reads were collapsed based on UMIs to generate a matrix of CellTag UMIs for each cell. Weighted Jaccard Similarity analysis was performed on data normalized for the total number of CellTags per cell, using the R package, *Proxy*. We opted to employ weighted analysis, based on observations that abundant CellTag expression is robust and stable between clones over the course of culture (**Fig.S4A**). This also places less emphasis on the less abundant CellTags which are more likely to arise due to noise, avoiding the need for any filtering based on expression levels. Lentiviral silencing has not presented an issue with this approach, where we can reliably detect CellTag expression to at least 11-weeks post labeling. Clonal relationships were visualized via a correlation matrix (**Fig.4A**) and clones were identified via hierarchical clustering as a group of 5 or more highly-related cells, based on CellTag overlap.

### CellTagging reveals extensive heterogeneity within each clone of reprogramming cells

iEPs self-renew *in vitro*, and can be passaged long-term^10,13^. **Figure 4B** demonstrates the emergence of clones over time, where over the first reprogramming timecourse we have detected 48 major cell clones. We selected these 48 clones and present them as a dendrogram, demonstrating that several clones have undergone significant expansion (**Fig.4C**). Of the benchmarked fibroblast population, which was split from the CellTagged population prior to reprogramming, only 2 clones were identified. In contrast, when we enriched for cells classifying as fully-reprogrammed iEPs (fractional iEP identity > 0.8, we could discern many clones, demonstrating their expansion as a result of their reprogramming (**Fig.S4B**). We were able to track the evolving identity of many clones over several timepoints, revealing the dynamics of reprogramming from cells descending from the same original fibroblast. Mapping these clones onto t-SNE shows that cells sharing clonal identity demonstrate a remarkable degree of heterogeneity in terms of reprogramming outcome (**Fig.4E**). In most instances, reprogramming cells derived from the same individual cell can occupy many different transcriptional states (**Fig.S4C**), suggesting that the same reprogrammed cell can adopt many different outcomes. By following individual clones, we were able to classify several broad reprogramming behaviors. For example, we see several instances where many descendants are located within early-transition and transition phases at timepoint 3, with some cells entering a fully-reprogrammed state (**Fig.4E; S4C**). By timepoint 4, though, many of these reprogrammed cells are lost with a concomitant gain in the ‘death’ cluster, suggesting that cells reprogram and then die. This could be described as a reprogramming ‘branchpoint’ toward a literal dead end. Alternatively, cells in the transition phase may begin to die and do not fully reprogram, and that the loss of fully reprogrammed cells arises due to reversion of these cells into the transition zone. This possibility is supported by the observation, in some instances, that the number of descendants in the transition zone increases (**Fig.S4C: clone B**). This could also be accounted for by relative expansion of these transition cells. Modified timing of CellTag delivery will help resolve these possibilities. Overall, this analysis demonstrates that reprogramming cells derived from the same initial cells, within a short timeframe can adopt many distinct transcriptional states. In agreement with previous reports, this apparent stochasticity would account for low reprogramming efficiency. In terms of overall efficiency, half of cells classified as iEPs are located within the major clones in the first timecourse. In the second timecourse, this is much more pronounced with 50% of all iEPs derived from the same single cell (**Fig.5A).** This suggests that many of the iEPs that were counted in previous colony formation assays (seeded one week after initiation of reprogramming) were derived from the same original cells, and therefore the actual efficiency of fibroblast to iEP reprogramming is much lower than the 1-2% originally reported^10,13^.

**Figure 5.**
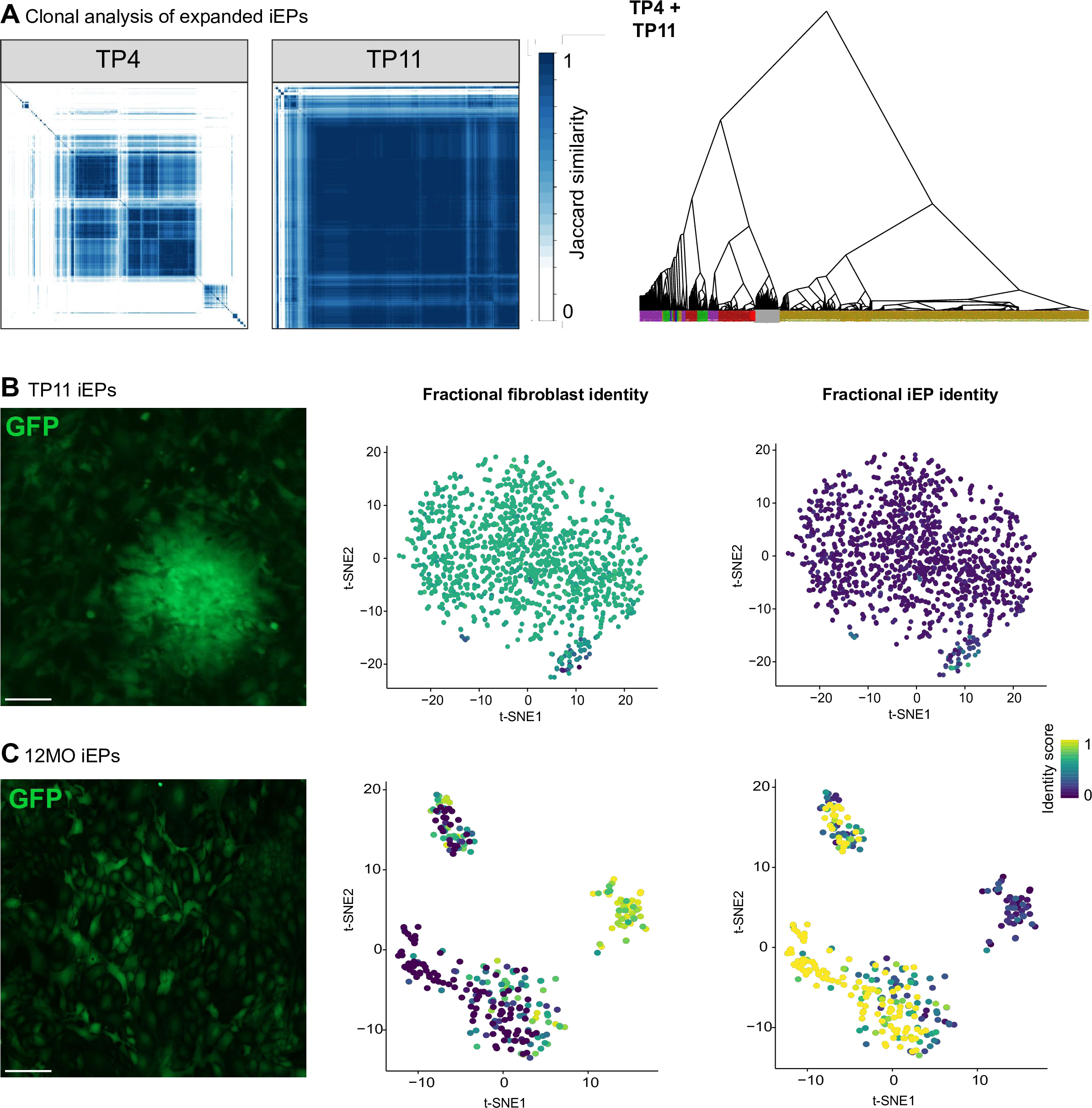
Clonal analysis of expanded iEPs. **(A)** Following reprogramming, iEPs can be expanded for long periods. In the second timecourse we continued passaging iEPs until 11-week post-initiation of reprogramming. Left panels: Clonal analysis of cells at timepoint 4 shows that a large proportion of cells are derived from the same clone. 50% of iEPs originate from the same fibroblast ancestor. Following culture until week 11, this clone has expanded to dominate the cell population. Right panel: dendrogram of all timepoint 4 and 11 cells from timecourse 2. **(B)** Image of iEPs at week 11: these expanded iEPs demonstrate phenotypic variability, with the emergence of small colonies of cells. Right panels: t-SNE visualizations of cell clusters and projection of fibroblast and iEP identity scores demonstrate that many fully reprogrammed cells have been lost from this population. Most cells of this population do not definitively score as either fibroblasts or iEPs. **(C)** Image of iEPs after 12 months of passaging: long-term expanded iEPs, unrelated to cells derived from timecourse 1 and 2. Right panels: t-SNE visualizations of cell clusters and projection of fibroblast and iEP identity scores demonstrate that cells resolve into three distinct clusters following long-term culture. iEPs are located in 2 clusters, whereas fibroblast identity can be clearly distinguished in a third, smaller cluster of cells. Scale bar = 100μM

### Fully-reprogrammed cells demonstrate long-term instability

At four weeks following initiation of reprogramming, we observe large clones of cells dominating the iEP populations (**Fig.4B;5A**). In the second timecourse we cultured these cells until week 11 and performed Drop-seq, followed by single-cell and clonal analysis. At week 4, cells derived from a single clone account for over 50% of the iEP population. Following passage for a further 5 weeks, over 90% of cells were derived from this one clone, originally stemming from an individual fibroblast (**Fig.5A**). Considering the self-renewal properties of this progenitor-like state, we expected most of these cells to classify as iEPs. Surprisingly, only a small fraction of the week 11 expanded iEP cells scored as iEPs (**Fig.5B;S3C**). Visual inspection of these expanded cells also revealed clear phenotypic heterogeneity within this population (**Fig.5B**). Together, these results suggest that fully reprogrammed iEPs may represent a transient state that is lost over time, perhaps to spontaneous differentiation or reversion to the original fibroblast state in culture. From differential expression analysis, no specific differentiation markers are identified within clusters, and fractional MEF identity scores moderately outside of the iEP cluster (**Fig.5B**). This suggests that iEPs in culture may revert back to a fibroblast-like state. This also supports our earlier observations that cells may revert back to a transition-like state during the active reprogramming phase. Even though the majority of these cells are derived from the same clone, is it possible that these sub-populations have arisen from distinct reprogramming trajectories. CellTagging of these stable lines will permit potential transitions of cells between these distinct transcriptional states to be monitored.

These findings raise questions about the long-term stability of the iEP reprogrammed state. To explore this in further detail, we performed Drop-seq profiling of an iEP cell line that had been continually passaged for one year. We had previously shown that this same line has the potential to functionally engraft a mouse model of acute colitis^13^. Surprisingly, many cells received high fractional iEP scores and were grouped within two subclusters, one of which expressed intestinal-associated genes *Reg3g* and *Gkn2.* A third cluster of cells expressing *Spp1* and *Mgp,* received high fractional fibroblast identity scores and may correspond to a reverted cell phenotype. Together, these results demonstrate that these established lines can undergo significant changes over time, and that dominance of the population by a small number of clones could significantly impact on the potential of each population as a whole.

## Discussion

Here, we present a combinatorial labeling method, CellTagging, which enables single-cell analysis of clonal dynamics during direct lineage reprogramming from fibroblasts to iEPs. Using Drop-seq, we have demonstrated that reprogramming to iEPs comprises distinct transition stages, suggesting roles for Igf2 and Notch signaling. Although many cells initiate reprogramming, full erasure of fibroblast identity and emergence of iEPs represents a rare event. Analyses of individual clones, originating from the same original cells, demonstrate a remarkable degree of heterogeneity with respect to reprogramming outcome. Longer-term tracking of fully-reprogrammed cells demonstrates that the expandable iEP population is derived from only a handful of initial fibroblasts. These resulting populations also display a great deal of heterogeneity, with evidence to suggest that iEPs may be in flux between distinct transcriptional states.

Originally, the intended target cell identity for iEPs was the hepatocyte^10^. Our previous analyses of these engineered cells revealed that intestinal identity was established in concert with hepatic fate, and that fibroblast identity remained intact^13^. These analyses were performed on microarray transcriptome data collected at the population level, leading to the conclusion that iEPs are partially reprogrammed. Single-cell analysis, presented here, reveals that the persistent fibroblast signatures in our previous studies were most likely derived from the sub-population of partially reprogrammed cells. Contrary to our previous findings, those rare cells that fully reprogram do appear to fully silence fibroblast identity. These observations demonstrate the utility of scRNA-seq and provide new insights into the reprogramming process.

Upon reprogramming to iEPs, a major group of cells (∼30%) initiate expression of iEP target genes. Although this initiation phase appears efficient, many cells continue to express fibroblast-associated genes in this transition period. Only those cells progressing to the fully reprogrammed state silence fibroblast gene expression programs. Amongst the direct lineage reprogramming strategies, generation of iEPs is among the most inefficient protocols with only 1-2% of fibroblasts generating iEPs^10,13^. This is in contrast to fibroblast to neuron direct reprogramming where 20% of fibroblasts yield induced neurons^22^. One method to generate neurons involves the expression of the TFs, Acsl1, Brn2, and Mytl1^20,22,36^. Recently, Mytl1 has been shown to be responsible for silencing non-neuronal identities to promote generation of neural fate in fibroblasts^37^. This raises the possibility that the Foxa1-Hnf4a reprogramming cocktail may be missing the Myt1l equivalent. If so, discovery of a third iEP reprogramming factor to silence all non-hepatic fate may increase the efficiency and fidelity of this particular engineering strategy.

In terms of reprogramming trajectories, our clonal tracing reveals that a single cell undergoing reprogramming does not follow a defined trajectory of transcriptional changes. Rather, descendants of the individual cell in which reprogramming was initiated may enter many asynchronous trajectories. Previous studies have demonstrated that reprogramming to pluripotency involves an initial stochastic phase^18,38^. The derivation of cell lines from an individual clone has demonstrated that all cells over time have the capacity to reprogram, and argue against the existence of ‘elite’ cells predisposed to reprogram^38^. Our results presented here are in agreement with these earlier findings, but the degree of heterogeneity derived from a single cell, under the same conditions, in short temporal order are surprising. It is tempting to speculate that this suggests that there are many more stochastic hurdles to overcome during cell fate conversion. One major barrier to reprogramming that has been proposed is the location of target genes ‘locked’ within heterochromatin^39^, and that access to these is a stochastic process. The clonal heterogeneity we observe may suggest that the accessibility of target genes fluctuates both over the short term, and at each step of the reprogramming process.

Although only a very small proportion of fibroblasts successfully reprogram to iEPs, these cells self-renew *in vitro* and can be expanded for long periods of time, perhaps indefinitely. One unexpected finding here is the dominance of an iEP population resulting from a single clone, after a relatively short period of expansion. Moreover, even though expanded iEPs were derived from only a handful of cells, the population exhibits clear cellular heterogeneity. In long-term cultured cells, iEPs appear to de-differentiate toward a fibroblast-like state, although iEP clusters are still maintained. CellTagging of these stable populations will reveal if the same individual cells can transition between these states, and the transcriptional control underlying this flux. In addition, our single-cell analyses have revealed a separate cluster of cells enriched for intestinal markers. This raises the possibility that these particular cells in our earlier intestinal transplantation studies were responsible for the functional engraftment we reported^13^. This population does not appear in every iEP line generated, suggesting that heterogeneity in reprogramming and the expansion of a small number of clones has a major impact on long-term cell potential. This is perhaps reflective of the skewed differentiation properties known to afflict many stem cell lines^40^.

In summary, here we have presented a method to facilitate analysis of clonal dynamics at single-cell resolution. This has revealed that in contrast to the initial heterogeneity of the reprogramming process, the resulting expanded cells lack clonal diversity. Beyond application to understanding the reprogramming process, the CellTagging approach will be broadly applicable to the study of any cell growth and differentiation process that is amenable to lentiviral transduction. The emergence of scRNA-seq approaches, such as the 10x Genomics Chromium system, that enable a higher single-cell capture rate will benefit this approach. In this case, smaller starting cell populations can be examined, which should help to increase capture of clones across multiple timepoints. In terms of temporal resolution, the CellTagging approach is easily adapted to multiple rounds of labeling to track clonal evolution in greater detail.

## Experimental Procedures

### Mice

Mouse Embryonic Fibroblasts were derived from the C57BL/6J strain (The Jackson laboratory: 000664). All animal procedures were based on animal care guidelines approved by the Institutional Animal Care and Use Committee.

### Generation of iEPs

Mouse embryonic fibroblasts were converted to iHeps/iEPs as in Sekiya and Suzuki (2011)^10^. Briefly, fibroblasts were prepared from E13.5 embryos and serially transduced with polyethylene glycol concentrated Hnf4a-t2a-Foxa1, followed by culture on gelatin for 2 weeks in hepato-medium (DMEM:F-12, supplemented with 10% FBS, 1 mg/ml insulin (Sigma), dexamethasone (Sigma-Aldrich), 10 mM nicotinamide (Sigma-Aldrich), 2 mM L-glutamine, 50 mM b-mercaptoethanol (Life Technologies), and penicillin/streptomycin, containing 20 ng/ml hepatocyte growth factor (Sigma-Aldrich), and 20 ng/ml epidermal growth factor (Sigma-Aldrich), after which the emerging iEPs were cultured on collagen.

### Lenti-and Retrovirus Production

Lentiviral particles were produced by transfecting 293T-17 cells (ATCC: CRL-11268) with pCMV-dR8.2 dvpr (Addgene plasmid 8455), and pCMV-VSVG (Addgene plasmid 8454). Virus was harvested 48 and 72 hr after transfection and PEG concentrated. Constructs were titered by serial dilution on 293T cells. Hnf4a-t2a-Foxa1 retrovirus was packaged with pCL-Eco (Imgenex), titered on fibroblasts, and cells transduced according to Sekiya and Suzuki^10^.

### CellTagging methodology

To generate CellTags, we introduced an 8nt variable region into the 3’UTR of GFP in the pSMAL lentiviral construct^41^ using a gBlock gene fragment (Integrated DNA Technologies) and megaprimer insertion. A complex library of CellTag constructs was used to generate lentivirus (above) which was then used to transduce fibroblasts at a multiplicity of infection of ∼10.

### Drop-seq procedure

Cells were dissociated using TrypLE Express (Gibco), washed in PBS containing 0.01% BSA and diluted to 100 cells/µl. These cells were processed by Drop-seq within 15 minutes of their harvest. Drop-seq was performed as previously described^23^. In brief, cells and beads were diluted to an estimated co-occupancy rate of 5% upon co-encapsulation. Two independent lots of beads (Macosko201110, ChemGenes Corporation, Wilmington MA) were used: 091615 (Timecourse 1), 032516B (Timecourse 2). Emulsions were collected and broken by perfluorooctanol (Sigma), followed by bead harvest and reverse transcription. After ExonucleaseI treatment, aliquots of 2,000 beads (∼ 100 cells) were amplified by PCR for 14 cycles. Following purification by addition of 0.6x AMPure XP beads (Agencourt), cDNA from an estimated 5,000 cells was tagmented by Nextera XT using 600 pg of cDNA input, as assessed by Tapestation (Agilent) analysis. Following further purification, 1ng of each library was used as input for GFP amplification for a further 12 cycles of PCR using the Drop-seq P5-TSO_Hybrid primer (AATGATACGGCGACCACCGAGATCTACACGCCTGTCCGCGGAAGCAGT GGTATCAACGCAGAGT*A*C) and a P7 hybrid primer complementary to the 3’ end of GFP (P7/Seq2/GFP Hybrid: CAAGCAGAAGACGGCATACGAGATGTGACTGGAGTTCAGACGTGTGCTCTTCCGATCTGGCATGGACGAGCTGTACAAGTAA). Following purification, the GFP amplicon was spiked into the main library at 5% of the total concentration. Libraries were sequenced on an Illumina HiSeq 2500 at 1.4pM, with custom priming (Read1CustSeqB Drop-seq primer).

### Computational methods Drop-seq alignment and digital gene expression matrix generation

After sequencing, scRNA-seq libraries were processed, filtered, and aligned as previously described^42^, including correction of barcode synthesis errors. This process, and the required tools, are further outlined online in the Drop-seq Alignment Cookbook (http://mccarrolllab.com/dropseq/). In an effort to facilitate downstream analyses the reference genome used during alignment was modified to include three transgenic sequences. The processed reads were aligned to the genome using STAR, default settings were used. Following alignment, digital gene expression (DGE) matrices were then generated for each timepoint from both time courses. Cells which expressed 200 or more genes were selected for inclusion in the matrices. For time course one this resulted in a combined 10,038 cells and 20,576 genes included in the digital expression matrix. The mean number of expressed genes in these cells was 1,215.908, and the mean UMI count was 2,636.718. This matrix was then filtered to include only cells with a UMI count of at least 1,000, cells in which the proportion of the UMI count attributable to mitochondrial genes was less than or equal to 10%, resulting in an expression matrix with 5,932 cells and 20,383 genes. This filtered expression matrix was used for all downstream analyses. The mean UMI count of the filtered matrix was 4120.815, and the mean number of expressed genes was 1,801.648. The same was performed for the second time course, resulting in a combined gene expression matrix of 8,335 cells and 21,023 genes. The mean UMI count and number of genes per cell, for the combined matrix was 5,502.507 and 1820.732 respectively. After filtering the matrix contained 5,414 cells and 20,934 genes, with a mean UMI count of 8,207.728 and a mean number of genes per cell of 2635.565. These filtered matrices were used for all downstream analysis.

### Cell cycle analysis and normalization

Following DGE filtering, cell cycle scores were generated for each cell and the data normalized. Cell cycle scores were generated as described in^43^, using the “pairs” based method. Briefly, this method compares the relative expression of gene pairs in a cell. Scores for each phase of the cell cycle are calculated based on the relative difference in expression of these genes. Finally, cells are assigned a phase of the cell cycle based upon the scores the cell receives for each individual phase. After calculating the cell cycle scores, the data was normalized using the “deconvolution” method described in^44^. This method pools cells and combines the expression values of the cells in a pool. The pooled expression values are used to calculate size-factors for normalization. These pool based normalization factors can then be “deconvoluted” into cell-specific normalization factors, which are then used to normalize each cell’s expression. This “deconvolution” normalization is an attempt to address the abundance of zero counts that is prevalent to scRNA-seq. The cell cycle scores and data normalization was facilitated by the Scater package, available on Bioconductor^45^.

### CellTag clonal analysis

Using the processed, filtered, and unmapped reads from an intermediate step of the alignment, reads that contained the CellTag “motif” were identified. From each of these reads, the UMI, Cell Barcode, and CellTag were extracted. For each cell barcode from the digital expression matrix, UMI counts were generated for each CellTag identified in the cell. The UMIs and CellTags were not collapsed based on hamming distance. Next, a matrix was constructed in which rows are Cell Barcodes and columns are CellTags. The value of any given location in the matrix, e.g. K x J, is equal to the UMI count of cell tag J, in cell K. The CellTag matrix was then filtered, removing CellTags appearing in >5% of cells in the first timepoint. Cells expressing more CellTags than two standard deviations from the mean were also removed, as these were likely to represent cell doublets. Weighted Jaccard analysis using the R package, Proxy was employed to calculate similarity between cells. Clones were classified as groups of 5 or more highly-related cells, visualized using the Corrplot package with hierarchical clustering.

### Seurat, Monocle, and quadratic programming analyses

After filtering and normalization of the DGE, the R package, Seurat^25^ was used to cluster and visualize the data. As the data was previously normalized, it was loaded into Seurat without normalization, scaling, or centering. Along with the expression data, meta data for each cell was included, containing information such as fractional fibroblast and iEP scores, cell cycle scores, cell cycle phase, and timepoint information. Seurat was used to remove unwanted variation from the gene expression, regressing out Timepoint, number of UMIs, proportion of mitochondrial UMIs, and cell cycle scores. After removing these unwanted sources of variation, highly variable genes were identified and used as input for dimensionality reduction via PCA. The resulting PCs and the correlated genes were examined to determine the number of components to include in downstream analysis. These principal components were then used as input to cluster the cells, using a graph-based approach. These clusters were visualized using tSNE. Unsupervised Monocle2^21,26^ analysis was used to order cells in pseudotime. Quadratic programming, previously described in^20^, was employed to score fibroblast and iEP identity. This approach was modified using bulk expression data collected previously^10^ and the KeyGenes^33^ platform to extract gene signatures of both cell types. The R Package, QuadProg was used for Quadratic Programming and generation of cell identity scores.

## Acknowledgements

We would like to thank members of the Morris lab for critical discussions, Rob Mitra for assistance with design of the GFP PCR amplification strategy, Steve McCarroll, Evan Macosko and Melissa Goldman for advice in establishing the Drop-seq platform, Barbara Treutlein for assistance with Quadratic Programming, John Dick for the kind gift of the pSMAL-GFP construct^41^, and Rich Head/Genome Technology Access Center for sequencing. This work was funded by the Children’s Discovery Institute of Washington University in St. Louis and St. Louis Children’s Hospital; and Washington University Digestive Diseases Research Core Center, National Institute of Diabetes and Digestive and Kidney Diseases [P30 DK052574].

